# To find better neural network models of human vision, find better neural network models of primate vision

**DOI:** 10.1101/688390

**Authors:** Kamila Maria Jozwik, Martin Schrimpf, Nancy Kanwisher, James J. DiCarlo

## Abstract

Specific deep artificial neural networks (ANNs) are the current best models of ventral visual processing and object recognition behavior in monkeys. We here explore whether models of non-human primate vision generalize to visual processing in the human primate brain. Specifically, we asked if model match to monkey IT is a predictor of model match to human IT, even when scoring those matches on different images. We found that the model match to monkey IT is a positive predictor of the model match to human IT (R = 0.36), and that this approach outperforms the current standard predictor of model accuracy on ImageNet. This suggests a more powerful approach for pre-selecting models as hypotheses of human brain processing.

## Introduction

In recent years, ANNs have revolutionized computer vision, achieving high performance on the object recognition ImageNet challenge. ImageNet performance gains have led to improvements in explaining primate ventral stream in monkeys (Yamins et al., 2013, Yamins et al., 2014) and humans (Khaligh-Razavi et al., 2014). Recently, high-performing models of object recognition and monkey recordings have become available (Schrimpf et al., 2018) but these have not yet been evaluated on human IT. Here we ask whether we can find a more powerful indicator of human IT predictivity across a range of state-of-the-art ANNs. Specifically, we tested whether ImageNet performance or model match to monkey IT (on different images) best predict model match to human IT.

## Methods

### Human fMRI

Human subjects (n=15, two separate groups) had their brain activity measured with a 3T fMRI scanner while they viewed 92 colored images of real-world objects segmented from their backgrounds and presented on a gray background (data from Cichy et al., 2014). Subjects performed a fixation task. Images were presented at the center of fixation (size: 2.9° visual angle, stimulus duration: 500 ms). A region of interest (IT) was defined in each subject using a mask consisting of bilateral fusiform and inferior temporal cortex (WFU Pickatlas, Maldjian et al., 2003).

### Monkey Neurophysiology

Neural responses to 2560 images (non-overlapping with the 92 human test images) were recorded from 168 IT neurons (data from Majaj et al., 2015). Recordings were acquired from two monkeys, each implanted with two Utah arrays in IT. Images were shown for 100ms at 8° visual angle. Neural firing rate was averaged in the window between 70 ms and 170 ms.

### Deep Artificial Neural Networks

We extracted activations from ANNs for the 2560 monkey test images and the 92 human test images using the Brain-Score platform (Schrimpf et al., 2018). All ANNs were trained to classify 1.2 million images into 1000 categories using the ImageNet dataset.

### Comparing Model Performance

We computed response-pattern dissimilarities between each pair of images and placed these in a representational dissimilarity matrix (RDM). We estimated model matches to human and monkey by correlating model (layers after each computation block) and data RDMs using Spearman correlation. We used the layer with the highest correlation with either monkey or human IT as “monkey IT” or “human IT” layer for each model. The “IT” layer selected for human and monkey could be different. We determined whether each of the model RDMs was significantly related to the human IT RDMs using a subject-as-random-effect analysis (one-sided Wilcoxon signed-rank test). We subsequently tested for differences in model performance between each pair of models using a subject-as-random-effect analysis (two-sided Wilcoxon signed-rank test). For each analysis, we corrected for multiple comparisons across models by controlling the FDR at 0.05. We correlated (Pearson R) monkey IT model match with human IT model match for the same set of ANNs.

### Control: ImageNet performance as an alternative predictor to find good models of human IT

Over the same set of 29 deep ANN models, we calculated the correlation (Pearson R) between ImageNet performance (top-1 error) with human IT predictivity.

## Results

### ImageNet performance vs monkey IT predictivity as a proxy for human IT predictivity

We tested whether ImageNet performance (top 1 error) is a good predictor of human IT predictivity. Across a set of 29 deep ANN models developed since ~2012, we plotted ImageNet performance vs human IT predictivity. We found a negative correlation (R = − 0.47, Figure 1), suggesting that ongoing improvements in ImageNet performance alone are unlikely to lead to better explanations of human IT. We then considered an alternative hypothesis that a way to find models of human visual processing is to find high performing models that are also better models of monkey vision. If this hypothesis is correct, then it must be robust to the choice of images – because the images are only used to select the models and all modern ANN models make internal response predictions for *any* image (i.e. they are all “image computable”). Thus, as a strong test of this hypothesis, we used different images to generate monkey and human ANN match scores. We correlated model match scores for monkey IT with model match scores for human IT for 29 ANNs. We found that indeed monkey IT match scores are positively correlated with human IT match scores (R = 0.36, Figure 2) suggesting that monkey IT predictivity is a better indicator of human IT predictivity than ImageNet performance.

**Figure 1.**
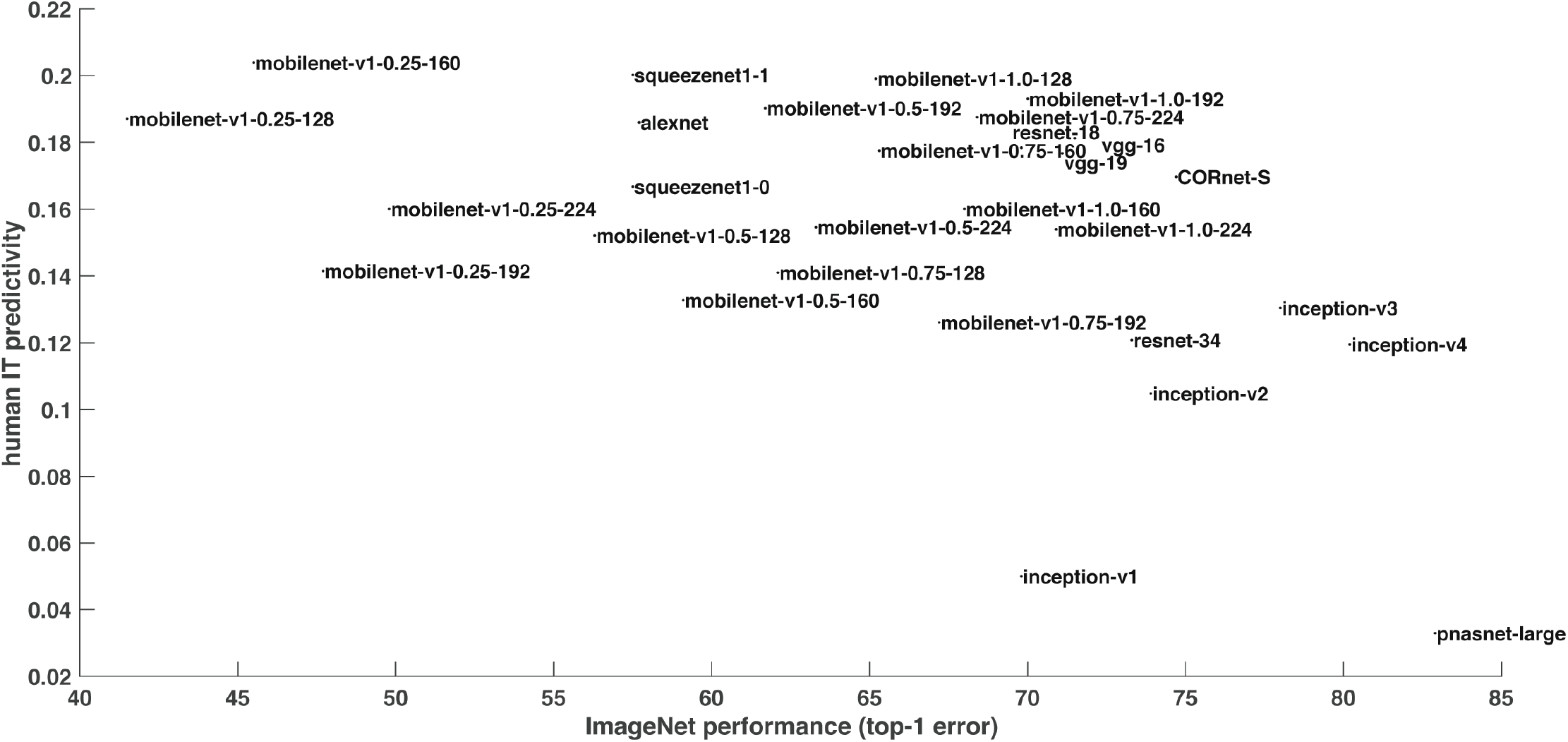
Correlation between ImageNet performance (top-1 error) and model match to human IT.

**Figure 2.**
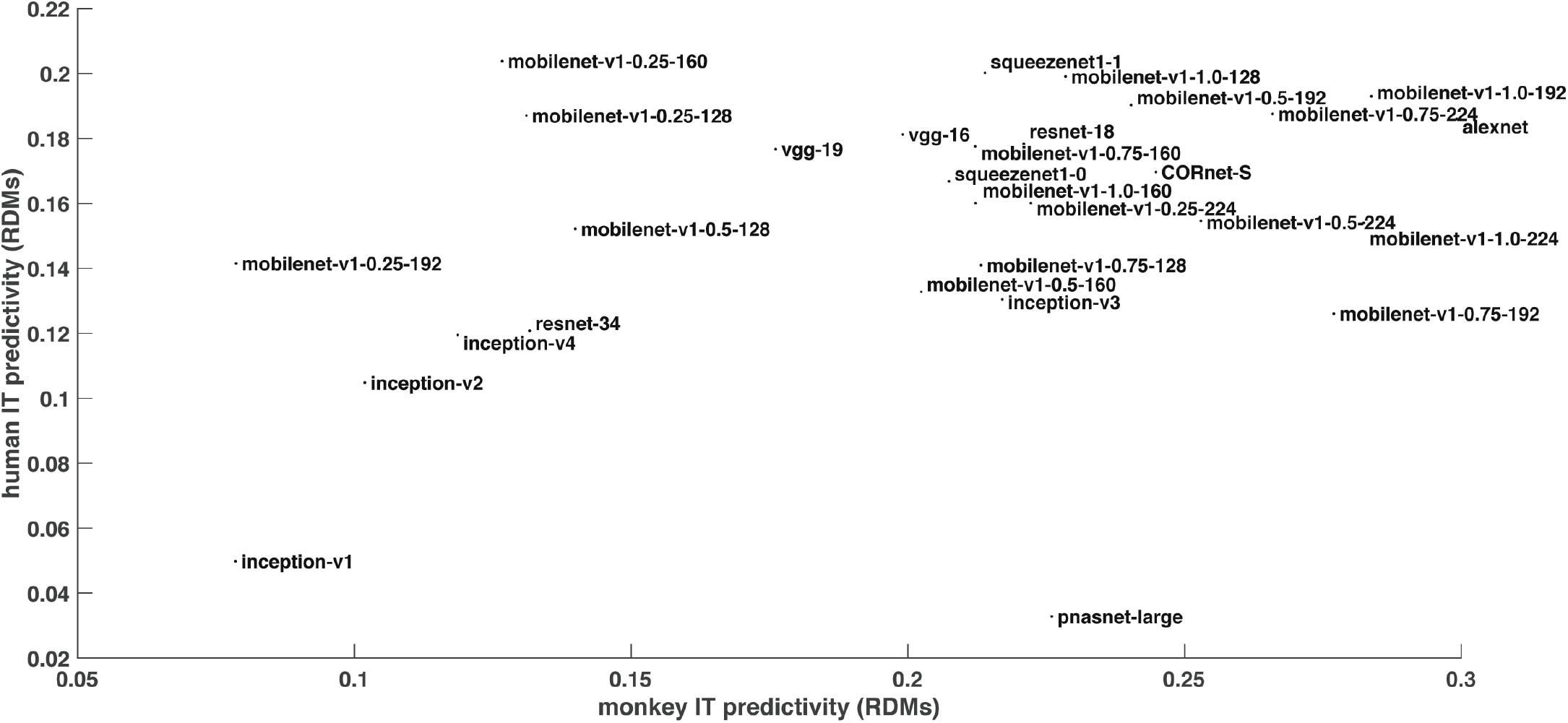
Correlation between model match to monkey IT and its match to human IT. The layers that best match IT could be different for human and monkey.

### Human IT predictivity

We assessed the ability of 29 ANNs to predict IT fMRI responses. Among the first ten models that have high IT predictivity are CORnets, squeezenet 1.1, some of the mobilenets, and alexnet (Figure 3). CORnets (Kubilius et al., 2018) have architectures that approximate the number and size of visual areas in the macaque brain. CORnet Z is feedforward, CORnet R and CORnet R2 are recurrent, and CORnet S is recurrent with skip connections. CORnet R has especially high human IT predictivity. CORnets have been shown to be one of the best models of the macaque brain. Here we show that CORnets are also good models for the human brain. We submitted the best layers of the best models (and their combinations) for the Algonauts challenge (Cichy et al., 2019), a competition aimed at finding models that best predict human fMRI and MEG brain representation.

**Figure 3.**
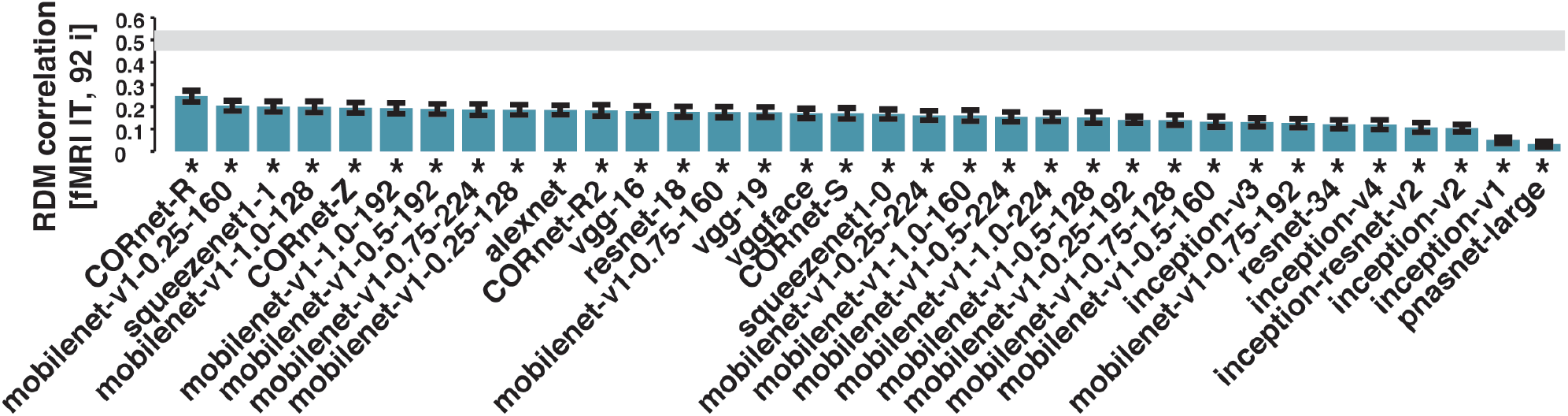
Predictivity of human IT responses. Bars show the correlation between the human IT RDM and each model RDM. A significant correlation between a model RDM and the IT RDM is indicated by an asterisk. We used subject-as-random-effect analyses for inference and error bars. The gray bar represents the noise ceiling, indicating the expected performance of the true model given the noise in the data.

## Discussion

We showed that a model’s match to monkey IT is a better predictor of its match to human IT than is the performance (accuracy) of the model on ImageNet. All ANNs tested here were optimized for ImageNet performance and reach high levels of top-1 error accuracy, but while ImageNet is extremely valuable for training parameters, accuracy levels alone seem to not predict human IT representations. It is possible that with a higher number of images and neuronal recordings from monkey IT a model match to human IT would be even better. This remains to be tested.

## Acknowledgments

This work was funded by the Sir Henry Wellcome Postdoctoral Fellowship (206521/Z/17/Z) to KMJ, the Semiconductor Research Corporation (SRC) to MS, the National Institutes of Health grant (DP1HD091947) to NK, the Simons Foundation grants (SCGB [325500, 542965]) to JJD, and the Center for Brains, Minds and Machines (NSF STC award CCF-1231216).

